# A simple and general approach to control the activity of DNA processing enzymes

**DOI:** 10.1101/2021.11.08.467724

**Authors:** Merve-Zeynep Kesici, Philip Tinnefeld, Andrés Manuel Vera

## Abstract

DNA processing enzymes, such as DNA polymerases and endonucleases, have found many applications in biotechnology, molecular diagnostics, and synthetic biology, among others. The development of enzymes with controllable activity, such as hot-start or light-activatable versions, has boosted their applications and improved the sensitivity and specificity of the existing ones. However, current approaches to produce controllable enzymes are experimentally demanding to develop and case specific. Here, we introduce a simple and general method to design light-start DNA processing enzymes. In order to prove its versatility, we applied our method to three DNA polymerases commonly used in biotechnology, including the Phi29 (mesophilic), Taq and Pfu polymerases, and one restriction enzyme. Light-start enzymes showed suppressed polymerase, exonuclease and endonuclease activity until they were re-activated by an UV pulse. Finally, we applied our enzymes to common molecular biology assays, and showed comparable performance to commercial hot-start enzymes.

## INTRODUCTION

DNA processing enzymes, such as DNA polymerases, nucleases and ligases, have been extremely useful tools in biotechnology, molecular biology, genetic engineering, molecular diagnostics methods and synthetic biology approaches (1–6). Over the last years, the development of enzymes with controllable activity has drawn attention as it allows minimizing secondary effects of undesired activity and triggering specific processes at will (7–11). One classical example is the development of hot-start PCR approaches (9,11–16). DNA polymerases are arguably one of the most successful applications of enzymes to biotechnology with applications in molecular biology, genome sequencing, sensing, and diagnostics among others (5,17). Yet, a common practical problem of these assays is that significant polymerase or proofreading activities (3’ to 5’ exonuclease activity) of the DNA polymerase during sample preparation can lead to loss of product yield and to lowered sensitivity and specificity (9–11,14,18,19). Polymerase activity can elongate missprimed events, including primer dimers and unspecific off target binding events, resulting in by-products that can compete with the desired amplicon. 3’ terminal primer degradation by 3’ to 5’ exonuclease activity can enhance off-target binding also inducing misspriming. Furthermore, exonucleolitic degradation of primers and template decrease the overall product yield (9–11,13,14,18,19). In hot-start, the reaction is blocked till the reaction mixture reaches an elevated, hot-start temperature (9,11,13,16,20). Other common strategy has been the design of photoactivatable enzymes, which has the additional advantage of allowing both space and time controlled enzymatic activity. Controlled PCR, DNA recombination and editing, optically controlled CRISPR-Cas9 genome editing, and the development of genetic switches has been possible thanks to photoactivatable enzyme versions (7,8,12,21–24).

The common drawback of all of these strategies is that they are not easily generalizable. Most common hot-start strategies employ specific aptamers or antibodies to selectively block the DNA polymerases. The development of such aptamers and antibodies require time-consuming screening of libraries and laborious optimization steps and after all are specific for a certain polymerase (9,15,16,20). Besides, the unavoidable heating step makes this type of strategy non-viable for mesophilic DNA polymerases, such as the Phi29 DNA polymerase (Phi29 pol), which is used for whole-genome amplification and sensing applications (5,25–27). The production of photoactivatable enzymes is commonly based in the incorporation of photosensitive unnatural amino acids and require detailed structural and mechanistic information of the enzyme (7,12,22–24). A meticulous study of the case is necessary to place the photosensitive residue in a position that blocks the activity in off state, and release it only after photoactivation. The latter requires robust knowledge of the enzyme and the devising a suitable strategy can be very challenging. Overall, these problems call for strategies of broader application and easier implementation to control the enzymatic activity of DNA processing enzymes. Here we present a quite general method to produce photoactivatable enzymes. We show the successful development of three light-activatable DNA polymerases and one nuclease, and prove their applicability to classical molecular biology methods.

## MATERIAL AND METHODS

### Construct design and molecular cloning

Binding of the ssDNA to the enzymes was achieved by site-specific incorporation of the unnatural amino acid 4-Azido-L-phenylalanine (azPhenylalanine). Using the amber codon suppression (TGA) strategy (28), this unnatural amino acid was included the sequence glycine-serine-azPhenylalanine-serine-glycine, which was added at the c-terminal. The SG-GS stretch provides conformational flexibility and accessibility, and site-specific labeling can be achieved without compromising any of the native residues of the enzymes. For the three DNA polymerases (N69D Phi29 pol (29), Pfu Pol and Taq pol) and the StuI restriction enzyme, codon-optimized genes for expression in *Escherichia coli* were designed using a web server (http://genomes.urv.es/OPTIMIZER/) and synthetic genes were purchased from Eurofins Genomics (Germany). The Pfu Pol, Taq pol and StuI genes were subcloned for expression in the pET24-d plasmid (Novagen, Merck Millipore, Germany) using its NcoI and XhoI restriction sites. The Phi29 pol gene was subcloned into pET21-a (Novagen) between the NdeI and XhoI sites. Standard protocols for molecular cloning were used (6) and the *E. coli* strain XL1blue was used for all cloning steps. Enzymes for cloning were purchased from NE Biolabs (MA, USA). The amino acid sequence of the proteins can be found in the Protein sequence section of the Supporting Material

### Protein production and purification

The *E. coli* Bl21 star (DE3) strain transformed with the plasmid pEVOL-pAzF (a gift from Prof. Peter Schultz, Addgene plasmid #31186) was used for protein expression. pEVOL-pAzF *E. coli* cells co-transformed with the expression plasmids were grown with vigorous shaking at 37 °C in LB medium with the adequate antibiotics until they reached an OD600 of ≈ 1. After that, the cells were pelleted, washed with M9 minimal medium, and resuspended in M9 minimal medium supplemented with 0.2 mg/mL 4-azido-L-phenylalanine (Hycultec GmbH, Germany), 0.02% arabinose, and antibiotics. The culture was incubated at 37 °C for one hour and finally induced with 1 mM IPTG overnight at 16°C. Cells were lysed and purified by nickel affinity chromatography as described elsewhere (30). In the case of Taq and Pfu pol samples the lysate was heated to 75 °C for 20 min after lysis to yield samples free of DNA contamination from *E. coli*. Phi29 pol, Pfu pol and StuI were further purified by cationic exchange in a HiScreen HP SP column (GE Healthcare), and Taq pol by anionic exchange in a HiScreen Q SP column (GE Healthcare). StuI was further purified by hydrophobic interaction chromatography in a HiTrap Phenyl HP column (GE Healthcare). The ion-exchange chromatography were performed as a linear gradient from 20 mM Tris, 100 mM NaCl, 0.2 mM EDTA pH 7.4 buffer to 20 mM Tris, 1M NaCl, 0.2 mM EDTA. The purity of the proteins was higher than 95% as assessed by coomassie staining in SDS-PAGE gels, with the exception of StuI. The concentration was estimated spectrophotometrically using the theoretical extinction coefficient at 280 nm.

### Protein-Oligo coupling

All oligonucleotides were purchased from Biomers GmbH (Germany). Two different types of photocleavable linkers, PC linker (1-(2-Nitrophenyl)-1,3-propanediol, https://www.biomers.net/en/Catalog/Modifications/PCLin/INTMOD) and PC BMN (1-(2-Nitrophenyl)-1,4-butanediol, https://www.biomers.net/en/Catalog/Modifications/PCLBM/INTMOD) were used, both yielding efficient cleaving. In order to prevent the degradation of the oligonucleotides by the exonuclease activity of the polymerases, the DBCO group was included in the 3’ in the case of Phi29 and Pfu pol (3’-5’ exo activity) and in the 5’ for Taq pol (5’-3’ exo activity). The DBCO group was included in the 3’ in the case of StuI. The PC linker was included before the terminal nucleotide bearing the DBCO site. Table S1 summarizes the details of the DBCO-modified oligos and the enzymes to which they were attached. For oligo-labeling, the proteins were incubated for at least 3 hours with the DBCO-modified oligos at a molar ratio of 1:1,3 or 1:1,5 in light-tight tubes at room temperature. Afterwards, the samples were purified by ionic exchange, using a cationic exchange column followed by an anionic exchange column. The different charges of free polymerases, polymerase-ssDNA, and free oligos allowed for efficient separation with this purification scheme. In the case of StuI, some residual free protein co-eluted always with the StuI-ssDNA, even after including an extra hydrophobic interaction chromatography step (see Fig S3 and discussion there). The Phi29 pol, Taq pol and StuI samples were stored in 10 mM Tris-HCl, 100 mM KCl, 0.1 mM EDTA, 1mM DTT, 0.5% Tween-20, 0.5 % IGEPAL CA-630 and 50 % glycerol. Pfu pol samples were stored in 25 mM Tris, 0.1 mM EDTA, 1mM DTT, 0.5% Tween-20, 0.5 % IGEPAL CA-630 and 50 % glycerol. All enzymes were stored at −20 °C.

### Determination of the concentration of the Enzyme-Oligo constructs

We estimated the concentration of the DNA Polymerase-oligo samples building calibration curves. For each polymerase-oligo construct the unmodified polymerase was mixed with the respective DBCO oligo at a molar ratio of 1:1, and the absorbance at 260 and 280 nm measured for several dilutions of this sample. As the concentration of the polymerase in the sample was known, we performed a linear fit of absorbance vs enzyme concentration. Then, the absorbance at 260 and 280 nm of the polymerase-oligo samples were used to interpolate their concentration using the linear fits. Finally, the mean value of the estimated concentration at 260 and 280 nm was used as the sample concentration. For StuI_PC_oligo, the concentration was estimated using the theoretical absorption coefficient of the oligo at 260 nm.

### Light activation

The UV pulses were applied using either a 45-watt 315 nm UV-Pad (Vilber, France) or a handheld 6-watt 365 nm lamp (Analytikjena GmbH, Germany). The samples were irradiated just before the polymerization or digestion reaction was started.

### Multiply-primed amplification experiments

We used multiply-primed amplification (18) to test the activity of the Phi29 pol samples. T7 blue plasmid (Novagen, Merck Millipore) and human genomic DNA (Roche Diagnostics, Germany) were used as templates for the experiments in Figure 2 and S1a, and Figure 3a and S1b respectively. Random hexamers (Thermofisher Scientific, USA) and the template were heated up at 95 °C for 3 min and incubated afterwards for 5 min on ice for annealing. Reactions took place at 30 °C in 50 mM Tris-HCl, 10 mM MgCl2, 10 mM (NH4)2SO4, 4 mM DTT, 0.02 % Tween20, 0.2 mg/ml BSA pH 7.5, supplemented with 0.5 mM dNTPs. Exonuclease protected hexamers (50 μM) were used for the plasmid amplification experiments, and unprotected hexamers (6.25 μM) for the failure-by-design ones. Template concentration was kept at 0.3 ng/μl in both cases. The enzymes were heat-inactivated for 15 min at 65 °C at the end of the experiments. To facilitate visualization of the reaction, the T7 blue amplified plasmid was digested with BamHI to linearize the concatenated plasmid copies. Quantification of the enzymatic activity (Figures 2d and 2e) was done using the fluorescence emission of SYBR I nucleic acid stain (Thermofisher Scientific). SYBR I was added to the samples after the reaction was completed and the fluorescence measured in a Real-time PCR machine (Rotor-Gene Q, Qiagen, USA). Triplicates for each conditions were measured and the mean value calculated. The fluorescence intensities were normalized to the higher fluorescence signal (the most active sample).

**Figure 1.**
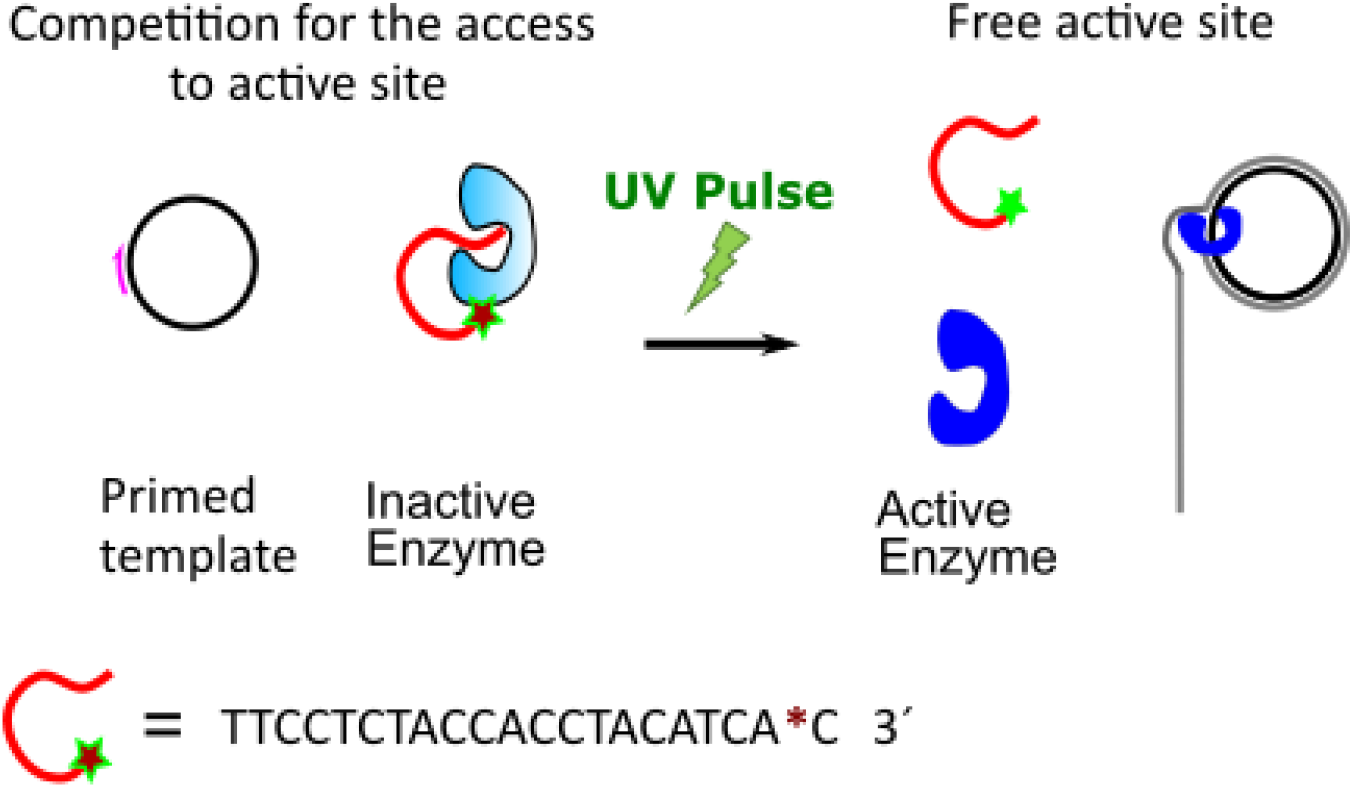
Schematics of light-start DNA processing enzymes. For illustration purposes, the case of a DNA polymerase is shown. The attachment of the ssDNA to the enzyme hampers the accessibility of the substrate to the active site leading to enzymatic blockage. Only after photocleavage of the bound DNA the activity is restored. The asterisk (*) denotes a photocleavable linker.

**Figure 2.**
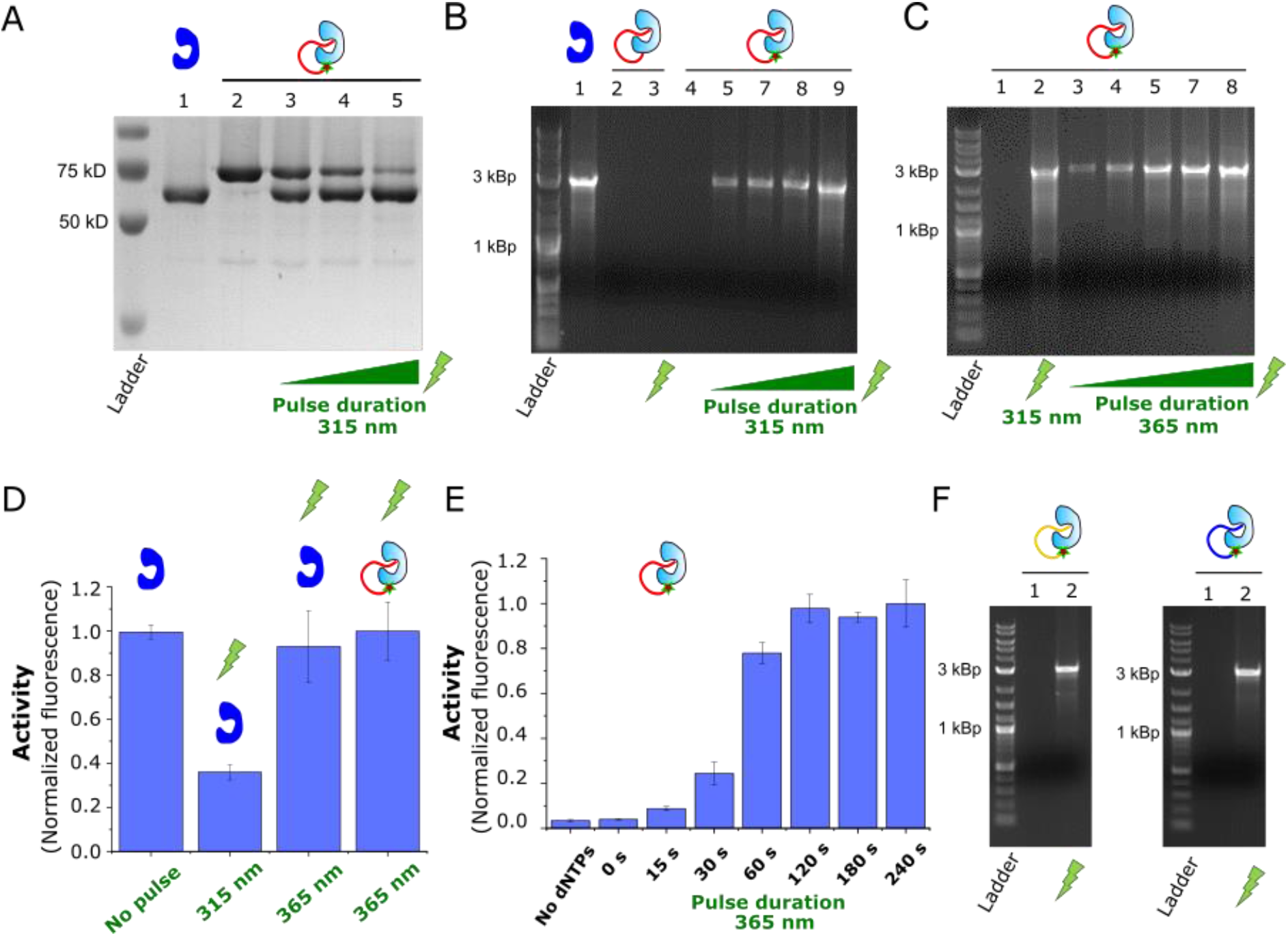
Light-activation of Phi29 DNA pol. **a)** SDS-Page showing the cleavage of the PC oligo from the Phi29 pol-PC_oligo construct by 315 nm UV light. Light pulses of increasing duration (5 s, 10 s and 20 s, lanes 2 to 5) produced a progressive fading of the Phi29 pol-PC_oligo band (lane 2 for comparison) correlated with the appearance and enrichment of the free enzyme band (lane 1 for comparison). **b)** Activity test in light-activated Phi29 pol-PC_oligo samples. Phi29 pol-PC_oligo samples illuminated with 315 nm UV pulses of increasing intensity and duration (1 s UV at 70% lamp intensity, 1 s, 2 s and 10 s, from lane 5 to 9 respectively) displayed amplified product of growing intensity. Activity was not observed in non-irradiated (lane 4) and Phi29 pol-oligo samples (lanes 2 and 3, 10 s 315 nm UV pulse was used for lane 3). **c)** Similar light-activated behaviour was observed with 365 nm UV light (from lane 3 to 8, light pulses of 5s, 10 s, 20 s, 30 s and 60 s respectively). For comparison, a sample illumined 10 s with 315 nm UV is included in lane 2. **d)** Effect of UV light in the assay. Unmodified enzyme was irradiated with 315 nm and 365 nm UV for 10 s and 120 s respectively. Non-illumined and irradiated Phi29 pol-PC_oligo samples (120 s 365 nm UV) are shown for comparison. **e)** Light activation curve of Phi29 pol-PC_oligo samples (365 nm UV). The signal in the non-illuminated samples (0 s) corresponds to fluorescence background, as it is not statistically different (p < 0.01) to inactive samples (no dNTPS). **f)** Light activation of Phi29 pol-PC_oligoScr (left gel) and Phi29 pol_oligo2 constructs (right gel). A light pulse of 365 nm 120 s was applied (lane 2). All activity assays were performed with 20 nM enzyme for 2 h at 30 °C. Phi29 pol is represented in blue and Phi29 pol-PC_Oligo in pale blue.

**Figure 3.**
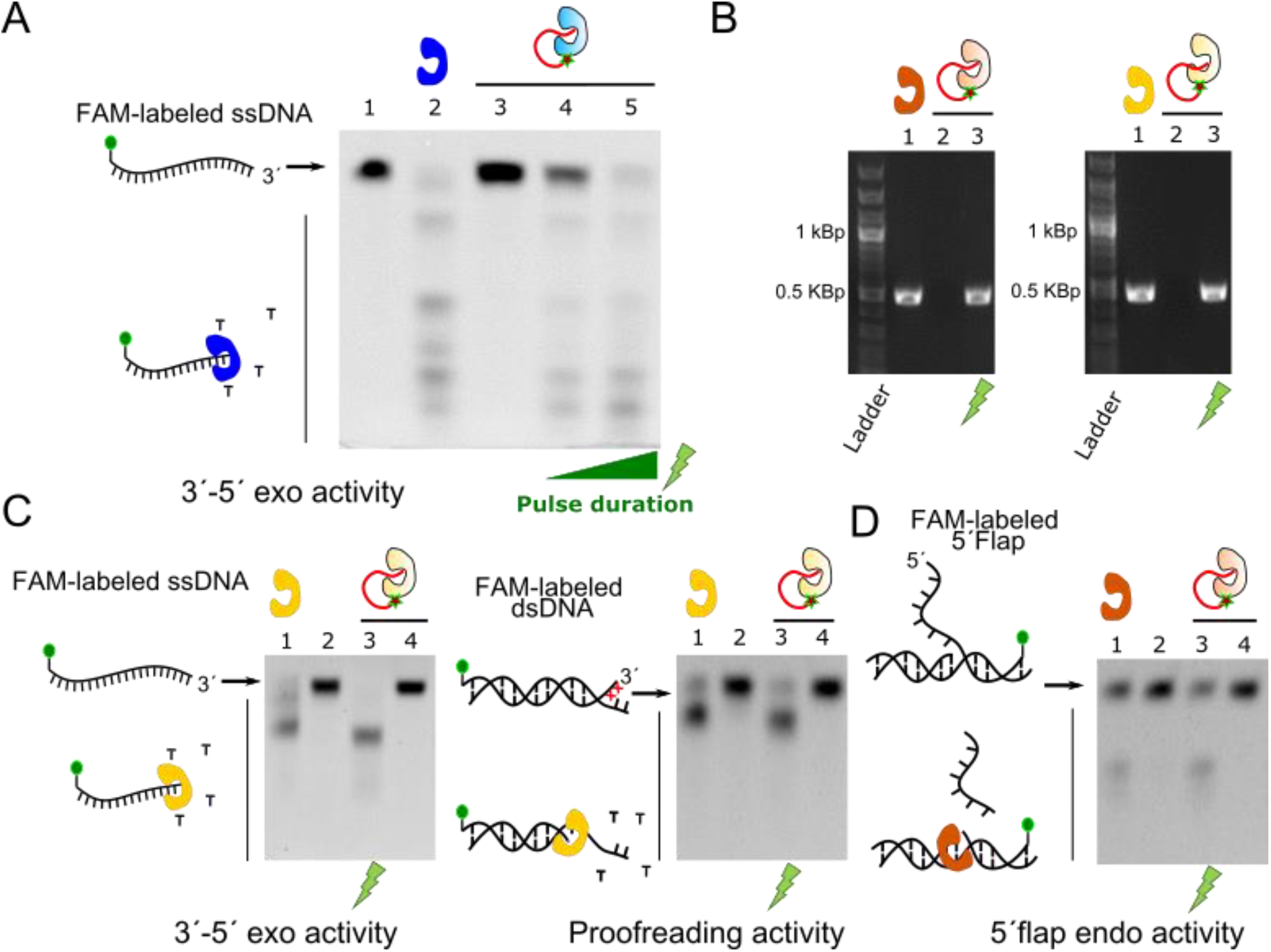
Polymerase and nuclease activity of different polymerases can be blocked. **a)** 3’ to 5’ exo activity of Phi29 pol-PC_oligo. Only light-activated samples (lane 4 and 5, 2 s and 10 s 315 nm pulses, respectively) recovered the exonuclease activity of the unmodified Phi29 pol (lane 2). **b)** PCR with Taq pol-PC_oligo (left gel) and Pfu pol-PC_oligo (right gel) samples. In both cases, PCR product was not detected in non-illuminated samples (lane 2). Only irradiated samples (lane 3) showed the PCR product present in the unmodified enzyme samples (lane 1). 20 nM of Taq pol and 15 nM Pfu pol were used. **c)** and **d)** Nuclease activity assays for Taq-PC_oligo and Pfu-PC_oligo. The exonucleolitic pattern of unmodified enzymes (lane1) was only recovered in light-activated samples (see lane 3 vs non-illuminated ones on lane 4). Left and right gel in **c)** show 3’ to 5’ exo and proofreading activities for Pfu pol-PC_oligo respectively. Gel in **d)** shows 5’ flap activity test for Taq-PC_oligo. Phi 29, Taq and Pfu pol are depicted in blue, brown, and yellow respectively, and the oligo-modified versions in the corresponding pale colors. A 10 s, 315 nm UV pulse was used for the experiments in **b)**, **c)** and **d)**.

### PCR experiments

PCRs were carried out in 50 mM Tris-HCl, 10 mM KCl, 10 mM (NH4)2SO4, 2mM MgSO4, 0.1 % Triton^®^ X-100, 0.1 mg/ml BSA pH 10.2 in a volume of 25 μl. As templates, 0.25 ng of a plasmid carrying a codon-optimized version of the human Cyclophilin A gene (PCRs in Figure 3b) and *E.coli* genomic DNA were used (PCRs in Figures 3b, 3c, S1c and S1d). *E.coli* genomic material was prepared from 5 ml overnight culture of XL1blue cells. The cells were pelleted, washed out twice with water, and resuspended in 200 μl of molecular biology grade water. Finally, they were heated at 99 °C for 5 min, centrifuged for 2 min at 15.000 g, and the supernatant with the chromosomic DNA transferred to a fresh tube. This chromosomic sample was diluted 10 times and 0.5 μl was used for the PCRs. Cycling parameter and primer sequences can be found in Table S2 and S3 respectively.

### Nuclease activity test

The test on 3’ to 5’ exonuclease activity of Phi29 samples were conducted using a 5’ FAM-labeled ssDNA oligonucleotide (FAM exo 3’ activity ssDNA, see table S4). 10 nM of enzyme was incubated with 50 nM ssDNA substrate for 10 min at 30 °C in 50 mM Tris-HCl, 10 mM MgCl2, 10 mM (NH4)2SO4, 4 mM DTT, 0.02 % Tween20, 0.2 mg/ml BSA pH 7.5. Exonuclease tests for Pfu samples were performed likewise but in 20 mM Tris-HCl, 10mM KCl, 10mM (NH4)2SO4, 2mM MgSO4, 0.1% Triton^®^ X-100 and 0.1mg/ml BSA pH 8.8 at 45°C for 10 minutes. The 3’ to 5’ exonuclease proofreading activity of Pfu was probed using a dye-labeled dsDNA with a five bases mismatch at the 3’ of the labeled oligonucleotide (FAM exo 3’ reverse + Exo 3’ mismatch forward, see table S4). The dye-labeled dsDNA was annealed forehand using a temperature gradient, and the reaction took place using 5 nM of enzyme and 50 nM of dsDNA at 45°C for 1 minute. The cleavage of single-stranded arms at the bifurcated end of base-paired duplexes by Taq pol (5’flap endonuclease activity) was used to test the nuclease activity of the enzyme (31). A bifurcated junction was used as substrate, and a primer located 4 bases upstream of the bifurcation point was included as it is known to promote the nuclease activity (31). The oligonucleotide forming the 5’ protruding strand was labeled at its 3’ with FAM and the fork-like structure assembled in a temperature gradient (Exo 5’ taq fork FAM + Exo 5’ taq template + Exo 5’ taq 4pb gap, see table S4). The reactions took place for 20 min at 45 °C using 50 nM of enzyme and 50 nM of the substrate in 20mM Tris-HCl, 10 mM KCl, 10 mM (NH4)2SO4, 2mM MgSO4, 0.1% Triton^®^ X-100 pH 8.8. All the reactions were stopped by adding formamide-loading buffer (6) and heating at 85 °C for 3 min. The samples were run in 10-12 % Urea-PAGE gels and the fluorescence signal read in a Fusion FX6 EDGE imaging system (Vilber).

The activity of the restriction endonuclease StuI was measured using a dsDNA probe bearing a FAM fluorophore and BMN-Q535 quencher at opposite *termini* (FAM StuI BMN-Q535 quencher + StuI reverse, see Table S4). A restriction site for StuI was included in the dsDNA and upon digestion an increase in fluorescence is expected. The reaction was performed at 37° C in Smart buffer (New England Biolabs, USA) in a Real-time PCR machine (Rotor-Gene Q, Qiagen, USA), using 1 nM of enzyme and 200nM of the dsDNA probe. The fluorescence signal was normalized to the highest intensity in the experiment.

### Failure-by-design experiments

Using a controlled experimental setup, the goal of these experiments is to show that undesired enzymatic activity during sample handling can be detrimental. The enzymatic concentration, pre-incubation times, and conditions were chosen to produce an experimental failure in case of significant exonuclease or polymerase activity during the pre-incubation step. Pre-incubations were carried out always at 25 °C to prove that the undesired activity would be present during sample handling as well. The enzymes were light-activated either at the end or at the beginning of the pre-incubation step. Experimental fail is expected in the latter, as the enzymes are active during the pre-incubation. In the case of Phi 29 pol-PC_oligo and Pfu pol-PC_oligo constructs, the failure-by-design experiments aimed at proving decreased amplification yield in case of undesired exonuclease degradation of the primers. For Taq pol-PC_oligo experiments, the primers were deliberately designed to form dimers at their 3’ ends. Polymerase activity would result in elongation during the pre-incubation step, and the elongated primer dimers would eventually compete with the desired PCR product. Pre-Incubation times were set to 2 hour, 1 hour and 20 min for the case of Phi29, Pfu and Taq pols experiments respectively (optimal incubation times were experimentally determined).

## RESULTS

### Photoactivatable Phi 29 polymerase

Serendipitously, we observed enzymatic inactivation of the Phi29 pol after binding a DNA oligonucleotide to the enzyme. We covalently attached the oligo to the enzyme using click chemistry (32), incorporating the unnatural amino acid 4-azido-L-phenylalanine (28) in the enzyme and reacting with a DBCO-modified oligonucleotide. As the unnatural amino acid was incorporated in an additional stretch of glycine-serine residues added by us at the *c-terminus* of the enzyme, we ruled out chemical modification of a key residue during the functionalization as the cause of the inactivation. We reasoned that having one of the natural substrate of the polymerase (ssDNA (33)) attached to it, could produce the effective blockage of the enzyme by means of binding to the protein’s cleft and competing for the accessibility to the active site (either obstructing the access to the active site or directly competing for it). In this case, the inactivation of the enzyme would be reversible, and controlled cleavage of the oligonucleotide would result in enzymatic reactivation (see Figure 1).

In order to test our hypothesis, we included an o-nitrobenzyl-based photocleavable (PC) linker (see experimental section for details) between the first and the second nucleotide proximal to the anchoring point of the enzyme (see Figure 1). In this configuration, a short UV pulse will release the oligonucleotide from the enzyme and reactivate it, giving us control over the activity. In Figure 1a, an SDS-PAGE gel shows the effect of UV pulses of different duration on the light-sensitive enzyme-oligo complex. A band corresponding to unmodified enzyme appeared in the irradiated samples (see lanes 3-5 in Figure 2a), and as expected the proportion of this population correlated positively with duration of the light pulse. Enzymatic reactivation was tested by multiply-primed rolling circle amplification of plasmidic DNA (18). Phi29 pol bound to a regular oligo (Phi29 pol-oligo) and non-irradiated Phi29 pol attached to a PC oligo (Phi29 pol-PC_oligo) showed no detectable activity after two hours of reaction (Figure 2b, lanes 2 and 4), as opposed to unmodified Phi29 pol enzyme, where an intense DNA band was observed in the agarose gel due to DNA polymerization (lane 1). This assures that enzymatic inhibition was also achieved for the PC variant. Furthermore, and more interestingly, the Phi29 pol-PC_oligo samples that were irradiated recovered the enzymatic activity, and this reactivation was correlated with the intensity and duration of the light pulse (Figure 2b, lanes 5-9). This reactivation was not observed in irradiated Phi29 pol_oligo (lane 3), confirming that the reactivation is specific to oligonucleotide cleavage.

We tested two different UV wavelengths to reactivate our enzymes, 315 nm and the less harmful 365 nm. Efficient enzymatic reactivation was observed for both wavelengths (Figure 2b and 2c), although shorter pulses seem to be required for the 315 nm wavelength (see Figure 2c, lane 2 vs lane 4). In order to calculate the reactivation efficiency, we first investigated any possible negative effects of the UV light in the reaction (including effect on the template, primers or enzyme). We studied the effect of both types of UV lights on the unmodified enzyme, as this allows us to decouple the reactivation efficiency from any negative effect caused by UV. We observed that short 10 s pulses with the 315 nm UV light produced a significant reduction of the activity, while 120 s pulses with the 365 nm UV did not have any negative effect on the reaction (see Figure 2d). Therefore, we used the 365 nm to estimate the reactivation efficiency. Under the same experimental conditions, the Phi29 pol-PC_oligo samples recovered the same level of activity as the wild type enzyme after a 120 s pulse with 365 nm UV light (Figure 2d). Furthermore, we confirmed that the Phi29 pol-PC_oligo enzyme reaches already saturation in the reactivation curve after 120 s (see Figure 2e). Altogether, these results suggest that the enzyme can recover full activity. Besides, we achieved a tight blockage of the enzymatic activity. We did not detect residual activity in the non-illuminated samples, as there was no statistical difference (p < 0.01) with samples that are not able to polymerize (samples without dNTPs, see Figure 2e). Even experiments performed at high concentration of the enzyme (150 nM) did not show activity in the non-illuminated samples (see Figure S1a), further confirming a severe inactivation of the enzyme.

In order to confirm that the blockage of enzymatic activity was mediated by unspecific obstruction by the coupled oligo and not by sequence-specific interactions, we used a scrambled version of the blocking oligonucleotide, which has the same nucleobase composition but in a random order (Phi29 pol-PC_oligoScr, see Supporting information for details). This Phi29 pol-PC_oligoScr version also inhibited activity that was recovered with a light pulse (Figure 2f, left). Moreover, to rule out any bias caused by the nucleotide composition, we blocked the enzyme with a third oligonucleotide with a completely different sequence (Phi29 pol_oligo2, Supporting information for details). This third type of modified Phi29 pol still displayed the same light-activated behaviour (Figure 2f, right).

### Blockade of the exonuclease activity

The proposed inhibition mechanism would also provide blockage of the 3’ to 5’ exonuclease (3’-5’ exo) activity of the Phi29 pol, as the oligonucleotide bound to the enzyme might also successfully compete with other exonuclease substrates provided it still hampers the access to the active site. We devised a test to characterize the 3’-5’ exo activity of our Phi29 pol constructs using a 5’ fluorophore-labeled (6-Carboxyfluorescein, 6-FAM) single stranded DNA probe (FAM-labeled ssDNA). Incubation of the labeled oligo with the unmodified Phi29 pol (Figure 3a, lane 2) showed a drastic drop in the intensity of the full-length FAM-labeled ssDNA and new populations of shortened FAM-labeled ssDNA when compared with the untreated sample (see lane 1 in Figure 3a). This pattern is also observed in the UV-activated Phi29 pol-PC_oligo samples, correlating again the degree of exonucleolitcc digestion of the FAM-labeled ssDNA with the duration of the light pulse (Figure 3a, lanes 4 and 5). On the contrary, when the Phi29 pol-PC_oligo sample is not activated by light, the FAM-labeled ssDNA remains intact (Fig 3a, lane 3). Altogether, these results demonstrate the controlled blockage of the 3’-5’ exo activity by the Phi29 pol-PC_oligo. Overall, our data confirm that our approach allows for the blockage and controlled reactivation of a thermolabile DNA polymerase, which is not possible with Hot-start approaches.

### Photoactivatable Taq and Pfu DNA polymerases

Our results with the Phi29 pol pointed to unspecific competition-based blockage of the enzyme by the covalently bound oligonucleotide. As the position of the modification was not rationally designed, we hypothesized that the same effect might be observed for other DNA polymerases as long as the oligonucleotide has enough flexibility to reach the active sites. Therefore, we implemented the same strategy in two other DNA polymerases widely used in biotechnology, the Taq and the Pfu DNA polymerases (for a discussion on how exonuclease degradation of the blocking oligo was avoided see supporting information, Protein-oligo coupling section). The Taq pol is the workhorse for PCR applications in all laboratories around the world and Pfu pol is a classical low error rate polymerase for application where high fidelity is desired (10,19,20). As in the case of Phi29 pol we incorporated 4-azido-L-phenylalanine in an extra stretch of C-terminal GS residues and covalently bound the same light-sensitive oligonucleotide. We first proved the blockage of the polymerization activity by PCR. Figure 2b shows that amplification is only detected in Taq pol-PC_Oligo and Pfu pol-PC_Oligo samples that had been treated with a UV pulse and in wild type enzymes (see Figure 3b, lanes 1 and 3, left gel for Taq pol and right for Pfu pol, respectively). Non-illuminated samples show no visible DNA band in the agarose gel (lane 2, Figure 3b). Furthermore, we do not have indication that the fidelity of the enzymes is significantly affected by the reactivation approach (see Figure S2).

Likewise, we tested for the inhibition of the nuclease activity of these polymerases. The Pfu pol has 3’ to 5’ exo activity, which includes proofreading activity (10,19) (3’ degradation in double stranded DNA, dsDNA, with 3’ terminal mismatches). We test the 3’ to 5’ exo activity of the Pfu pol constructs in two types of fluorescently labeled substrates, including ssDNA (FAM-labeled ssDNA) and dsDNA with mismatches (FAM-labeled dsDNA, as a substrate for proofreading activity). In both cases, the Pfu pol-PC_oligo samples that were not photo-activated did not show visible degradation of the substrates (Figure 3c, see lanes 2 and 4, left and right gel for the FAM-labeled ssDNA and FAM-labeled dsDNA substrates, respectively). Only the light-activated samples showed the degradation pattern typical of the wild-type enzymes (see lanes 1 and 3 in Figure 3c for both substrates). Taq DNA pol possesses 5’ nuclease activity, including 5’ to 3’ exonuclease and 5’ flap nuclease activity (10,31). We test the 5’ flap nuclease activity of our samples as it provides a convenient way to detect the 5’ nuclease activity (Figure 3d). Similarly, the nuclease activity of the Taq pol-PC_oligo was inhibited until the samples were photo-activated (Figure 3d, see lanes 3 and 4 and comparison with the wild type enzyme in lane 1). Altogether, our results show that not only the polymerase activity was blocked in the enzymes, but also the nuclease activity.

### Generalizing the approach

Our data demonstrated that our approach to block the activity of enzymes works robustly in DNA polymerases. Next, we aimed to check the general applicability our strategy to other DNA processing enzymes. Specifically, we focus in devising a light-activated version of the StuI Type II restriction enzyme. Since our approach relies on binding an ssDNA to compete for the access to the active site, the application to StuI would be a very stringent test to our strategy, since StuI is a sequence-specific nuclease acting in dsDNA. Thus, we attached the same oligonucleotide used for the three DNA polymerases to the *c-terminus* of the StuI enzyme, and checked its activity on a quencher-fluorophore dsDNA bearing a StuI restriction site. The results, shown in Figure 4, showed an acute inhibition of the nuclease activity of the StuI-PC_oligo sample, which was recovered in the irradiated sample (see green symbols *vs* red symbols in Figure 4 for illuminated and non-illuminated samples respectively). Although the inhibition of the activity seems not to be as acute as in the case of the DNA polymerases, there is a remarkable blockage of the enzymatic activity, especially considering that we could not manage to purify oligo-modified enzyme free of unmodified protein (see the significant amount of unmodified StuI enzyme in gel in Figure S3). Canonical type II restriction enzymes are dimeric (34) and we interpreted the constant co-elution of both species after several chromatographic purification step as hybrid dimers. This could explain the severe inhibition achieved, as the unmodified enzyme might still be forming hybrid oligomers with oligo-bearing ones. Our results suggest that even though dsDNA is the canonical substrate for restriction enzymes, the enzyme may retain substantial binding affinity to single-stranded DNA, which mediates the blockage. Alternatively, steric hindrance by the oligonucleotide bound to the enzyme could also render the observed inhibition.

**Figure 4.**
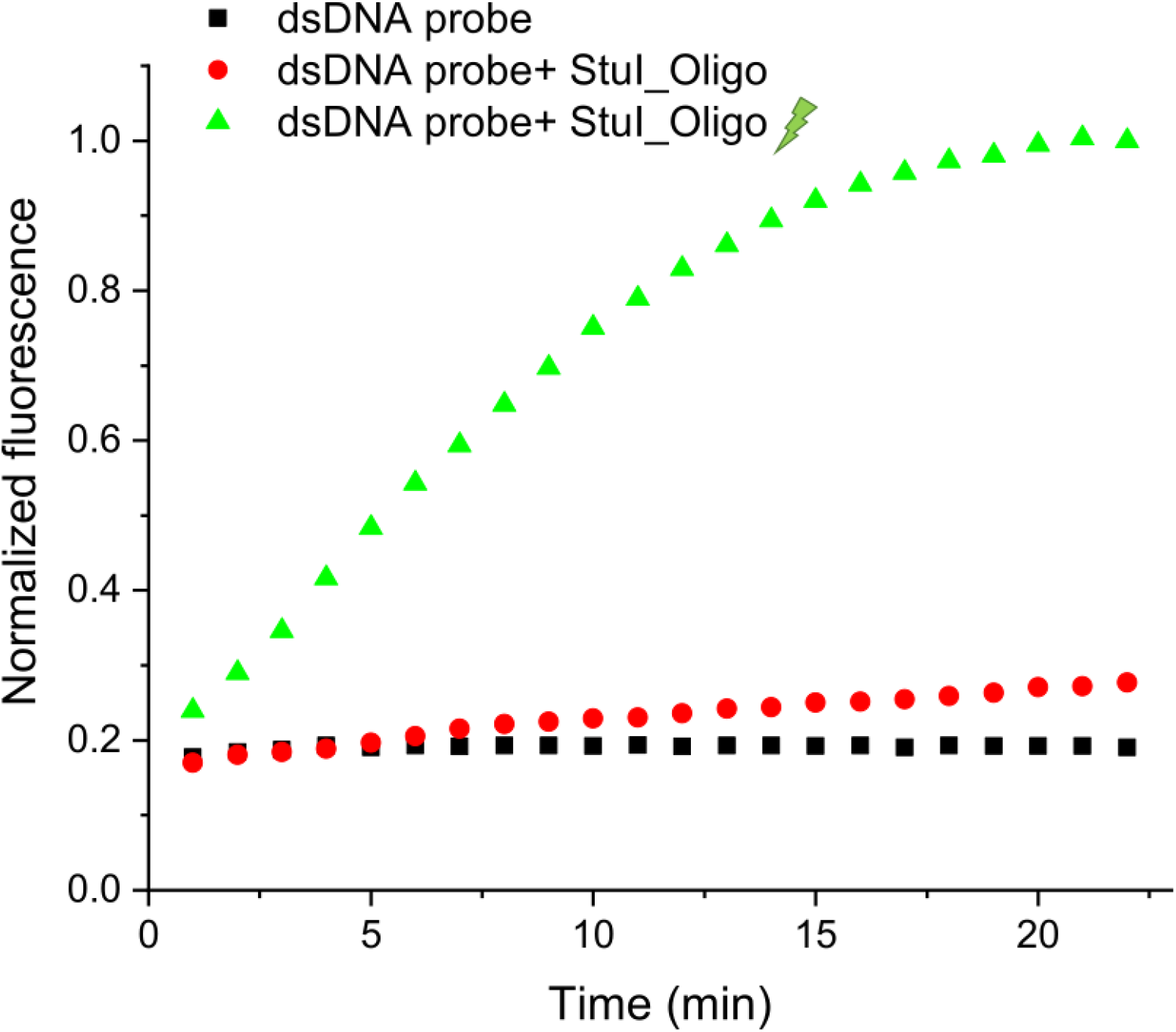
Selective activation of the restriction enzyme StuI. Activity assay for Stul enzyme. The irradiated Stul-PC_oligo sample (green symbols) showed a fast increase of the fluorescence signal, consistent with efficient digestion of the probe. This behaviour was largely supressed in non-illuminated samples (red symbols), where just a mild increase in the fluorescence is observed when compared with dsDNA probe alone (black symbols) .A light pulse of 120 s with 365nm wavelength UV light was used.

Altogether, our data suggests that the presented approach for controlled activation of DNA processing enzymes is of broad applicability and potentially transferable to many other enzymes

### Application of the light-activatable enzymes to classical biotechnological methods

Finally, as a proof of the relevance of our approach, we tested the applicability of our light-start DNA polymerases to common molecular biology applications. We designed a series of failure-by-design experiments, typically used to prove the goodness of hot-start approaches (13). Samples were preincubated before the assay and enzymatic activity during the incubation is expected to produce a detrimental effect (14,35). Specifically, samples were light-started at the beginning or the end of the pre-incubation and the negative effect was expected in the former, as the enzyme is active during the incubation.

Primer and template degradation by exonuclease activity is an issue in Phi29 pol and Pfu pol applications, producing decreased amplification yield in both cases and promoting unspecific off-target amplification in Pfu pol PCRs (10,18,19,26). We performed whole human genome amplification (25) using the Phi pol-PC_oligo enzyme to test for exonuclease protection. We observed a severe reduction of the product yield in the samples activated at the start of the incubation step (see comparison with post-incubation activation, Figure 5a and S1b). The same bias was observed for Pfu pol-PC_oligo, when we performed a PCR to amplify the Biotin ligase gene (Bir A) from *E. coli* chromosomic DNA (Figure 5b and S1c). These results are compatible with exonuclease degradation of the primers during the incubation and show the protection provided by our light-start enzymes.

**Figure 5.**
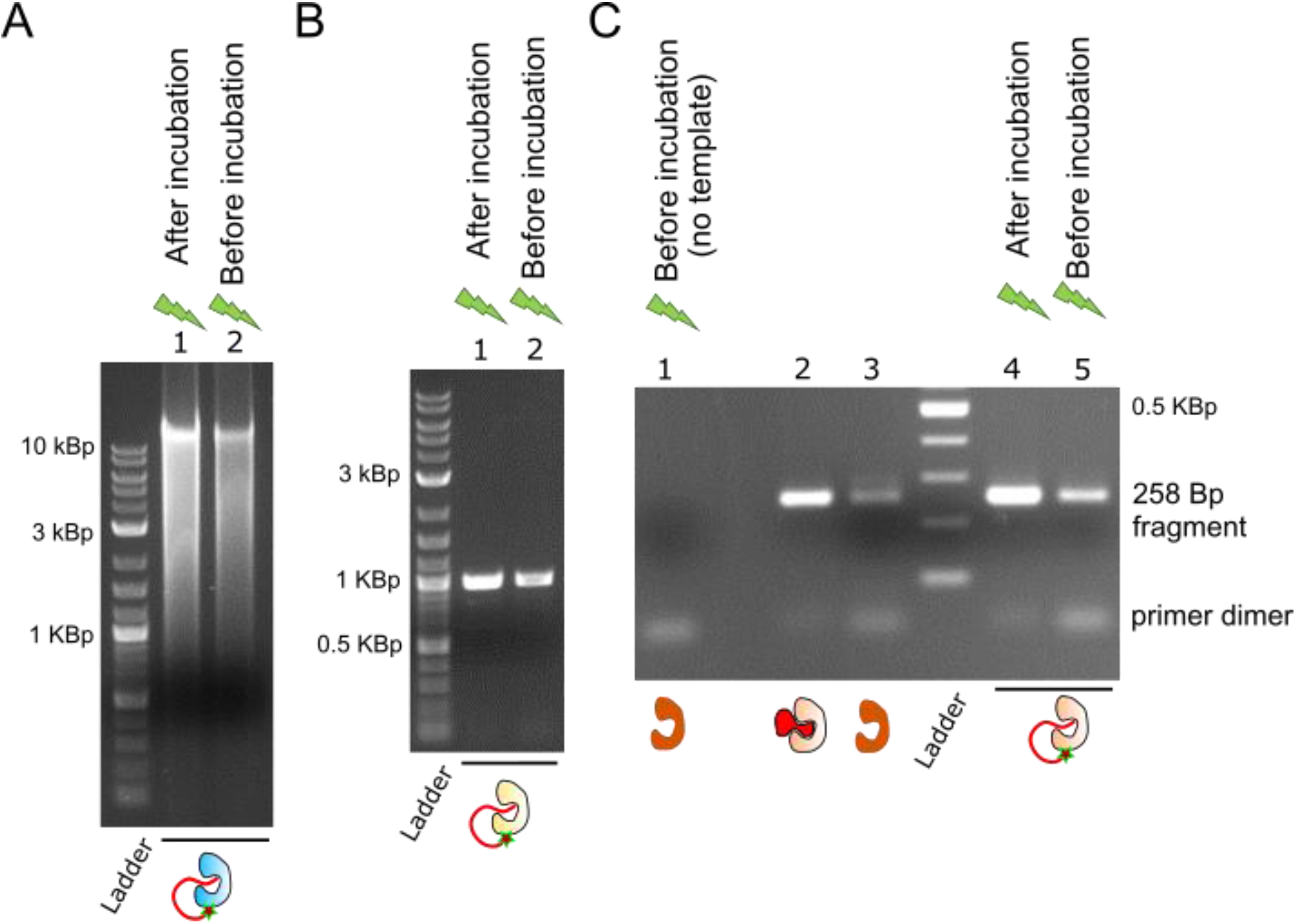
Light-start applications. **a)**, **b)** and **c)** failure-by-design experiments for Phi29 pol-PC_oligo, Pfu pol-PC_oligo, and Taq pol-PC_oligo, respectively. **a)** Whole genome amplification of human DNA by Phi29 pol-PC_oligo, and **b)** PCR amplification of *E. coli* Bir A gene by Pfu pol-PC_oligo. The samples that were kept inactive during the pre-incubation step showed increased yield of amplified products (lane 1), consistent with reduced exonuclease degradation of the primers. **c)** Light-start PCR shows protection against formation of primer dimers (lane 4 *vs* lane 5, see lane 1 for dimers reference) similar to that achieved by commercial hot-start Taq pol enzymes (lanes 2 and 3, NE Biolabs aptamer-based hot-start and standard Taq pol). Lane 1 shows a PCR performed without DNA template as reference for primer dimers formation. Enzymes concentrations were 120 nM, 7.5 nM and 50 nM for Phi29, Pfu and Taq samples. See Figure S1 for additional independent experiments.

In the case of Taq pol, elongation of missprimed primer during sample handling can lead to a loss of specificity in PCR (9,20). We PCR-amplified a 258 base pairs (Bp) fragment of *E. coli*’s Bir A gene and faulty designed the forward and reverse primers to anneal between their 3’ ends. These primer dimers if elongated during the incubation would compete with the PCR fragment. We observed that, while Taq pol-PC_oligo samples that were activated after the incubation showed a robust amplification of the 258 Bp fragment, in the samples photo-activated before the incubation the primers dimers competed with the desired fragment and produced an acute reduction in the PCR yield (see Figure 5c, lanes 1, 4 and 5, and Figure S1d). Furthermore, similar behaviour was observed using a commercial hot-start Taq polymerase based on aptamer blockage (Figure 5c, lanes 2 and 3).

Overall, we have proven that light-start polymerases are a robust alternative to traditional hot-start approaches. Unlike the latter, our approach can be generalized to thermolabile enzymes and we have shown that, unlike previous strategies (9,12,16,20,36), it is straightforward to implement and potentially applicable to diverse DNA polymerases.

## DISCUSSION

The application of the presented methodology to four different DNA processing enzymes, with diverse type of enzymatic activity, including DNA polymerase, exonuclease (5’-3’ and 3’-5’ exonuclease) and endonuclease activity shows promise as a general method to control the activity of these enzymes. In addition, producing the light-activatable version of the enzymes prove to be straightforward, as it sufficed to bind the photocleavable oligonucleotide in an extra stretch of aminoacids at the *c-teminus* of the enzyme. The latter is of particular practical interest because no specific design or knowledge of the enzymes needs to be taken into account. We are confident that this approach offers an alternative option before engaging in more complex and demanding strategies. Furthermore, the strategy would be potentially transferable to enzymes with strict dependence on dsDNA, by means of including a stem-loop structure in the blocking oligo.

## AVAILABILITY

All data and materials are available from the corresponding authors on request.

## SUPPLEMENTARY DATA

Supplementary Data are available at NAR online.

