## Supplementary material for "A simple and general approach to control the activity of DNA processing enzymes"

### *Supporting Information*

| Oligonucleotide | DNA polymerase-Oligo |
|---|---|
| ttcctctaccacctacatca*c-DBCO | Phi29 pol-PC_oligo/ Pfu pol-PC_oligo/StuI_PC_oligo |
| DBCO-t*tcctctaccacctacatcac | Taq pol-PC_oligo |
| cttcatcacactccatctcc*a-DBCO | Phi29 pol-PC_oligoScr |
| gcattacgtttggtggaccc*t-DBCO | Phi29 pol-PC_oligo2 |
| ttcctctaccacctacatcac-DBCO | Phi29 pol-oligo |

**Table S1. Sequence for the DBCO modified** oligonucleotides. The asterisk (*) indicates the position of the photocleavable linker.

| | N° cycles | Denaturing | Annealing | Extension |
|---|---|---|---|---|
| Cyclophilin A | 36 | 15 s at 95 °C | 20 s at 50 °C | 30s at 72 °C |
| Bir A gene (Pfu) | 39 | 15 s at 95 °C | 20 s at 56 °C | 1min:30s at 72 °C |
| Bir A gene (Taq) | 39 | 15 s at 95 °C | 20 s at 63 °C | 1min:30s at 72 °C |
| Bir A fragment | 39 | 15 s at 95 °C | 20 s at 59 °C | 30s at 72 °C |

**Table S2. Cycling parameters used for PCR experiments.** All PCR experiments start with an initial denaturing step (5 min, 95 °C), followed by the cycling loop, and a final 10 min elongation step at 72 °C. The different cycling parameters are summarized in this table and the primers in table S3.

| | Forward Primer | Reverse Primer |
|---|---|---|
| Cyclophilin A | TTCGCCATGGTTAACCCGACCGTTTTCTTCG | GAAGCTCGAGCTGACCGCAGTCCGCGATGG |
| Bir A gene | TCTACCATGGGCAAGGATAACACCGTGCCACTG | ATTAGCTCGAGTTTTTCTGCACTACGCAGGGA |
| Bir A fragment | AGAGTGTCGTTAATCAGGG | GCTCAAGTAATAAAGCCC |

**Table S3. Primers used for PCR**

| **ssDNA name** | **sequence** |
|---|---|
| FAM exo 3' activity ssDNA | FAM-tctctctctctctcctcttcctatattccgtacttc |
| FAM exo 3' reverse | FAM- gaagtacggaatataggaagaggagag |
| Exo 3' mismatch forward | tctctctctaatcgctcttcctatattccgtacttc |
| Exo 5' taq 4pb gap | ggatgagataggatgaagtacgg |
| Exo 5' taq template | tctctctctaatcgctcttcctatattccgtacttcatcctatctcatcc |
| Exo 5' taq fork FAM | ttacttcctaggaagagcgattagagagaga-FAM |
| FAM StuI BMN-Q535 quencher | FAM-ttcaggcctttt- Q535 |
| StuI reverse | aaaaggcctgaa |

**Table S4. Oligonucleotides used in the nuclease activity experiments.** FAM stands for 6-carboxyfluorescein.

### ***Protein sequences***

The position of the 4-Azido-L-phenylalanine incorporation is highlighted.

*Phi29 pol*

MPRKMYSCDFETTTKVEDCRVWAYGYMNIEDHSEYKIGNSLDEFMAWVLKVQADLYFHdLKFDGAFIINWLERNGFKWSA
DGLPNTYNTIISRMGQWYMIDICLGYKGKRKIHTVIYDSLKKLPFPVKKIAKDFKLTVLKGDIDYHKERPVGYKITPEEY
AYIKNDIQIIAEALLIQFKQGLDRMTAGSDSLKGFKDIITTKKFKKVFPTLSLGLDKEVRYAYRGGFTWLNDRFKEKEIG
EGMVFDVNSLYPAQMYSRLLPYGEPIVFEGKYVWDEDYPLHIQHIRCEFELKEGYIPTIQIKRSRFYKGNEYLKSSGGEI
ADLWLSNVDLELMKEHYDLYNVEYISGLKFKATTGLFKDFIDKWTYIKTTSEGAIKQLAKLMLNSLYGKFASNPDVTGKV
PYLKENGALGFRLGEEETKDPVYTPMGVFITAWARYTTITAAQACYDRIIYCDTDSIHLTGTEIPDVIKDIVDPKKLGYW
AHESTFKRAKYLRQKTYIQDIYMKEVDGKLVEGSPDDYTDIKFSVKCAGMTDKIKKEVTFENFKVGFSRKMKPKPVQVPG
GVVLVDDTFTIKSGazFGSLEHHHHHH

*Taq pol*

MGRGMLPLFEPKGRVLLVDGHHLAYRTFHALKGLTTSRGEPVQAVYGFAKSLLKALKEDGDAVIVVFDAKAPSFRHEAYGGYKAGR
APTPEDFPRQLALIKELVDLLGLARLEVPGYEADDVLASLAKKAEKEGYEVRILTADKDLYQLLSDRIHVLHPEGYLITPAWLWEK
YGLRPDQWADYRALTGDESDNLPGVKGIGEKTARKLLEEWGSLEALLKNLDRLKPAIREKILAHMDDLKLSWDLAKVRTDLPLEVD
FAKRREPDRERLRAFLERLEFGSLLHEFGLLESPKALEEAPWPPPEGAFVGFVLSRKEPMWADLLALAAARGGRVHRAPEPYKALR
DLKEARGLLAKDLSVLALREGLGLPPGDDPMLLAYLLDPSNTTPEGVARRYGGEWTEEAGERAALSERLFANLWGRLEGEERLLWL
YREVERPLSAVLAHMEATGVRLDVAYLRALSLEVAEEIARLEAEVFRLAGHPFNLNSRDQLERVLFDELGLPAIGKTEKTGKRSTS
AAVLEALREAHPIVEKILQYRELTKLKSTYIDPLPDLIHPRTGRLHTRFNQTATATGRLSSSDPNLQNIPVRTPLGQRIRRAFIAE
EGWLLVALDYSQIELRVLAHLSGDENLIRVFQEGRDIHTETASWMFGVPREAVDPLMRRAAKTINFGVLYGMSAHRLSQELAIPYE
EAQAFIERYFQSFPKVRAWIEKTLEEGRRRGYVETLFGRRRYVPDLEARVKSVREAAERMAFNMPVQGTAADLMKLAMVKLFPRLE
EMGARMLLQVHDELVLEAPKERAEAVARLAKEVMEGVYPLAVPLEVEVGIGEDWLSAKESGazFGSLEHHHHHH

*Pfu pol*

MGILDVDYITEEGKPVIRLFKKENGKFKIEHDRTFRPYIYALLRDDSKIEEVKKITGERHGKIVRIVDVEKVEKKFLGKPITVWKL
YLEHPQDVPTIREKVREHPAVVDIFEYDIPFAKRYLIDKGLIPMEGEEELKILAFDIETLYHEGEEFGKGPIIMISYADENEAKVI
TWKNIDLPYVEVVSSEREMIKRFLRIIREKDPDIIVTYNGDSFDFPYLAKRAEKLGIKLTIGRDGSEPKMQRIGDMTAVEVKGRIH
FDLYHVITRTINLPTYTLEAVYEAIFGKPKEKVYADEIAKAWESGENLERVAKYSMEDAKATYELGKEFLPMEIQLSRLVGQPLWD
VSRSSTGNLVEWFLLRKAYERNEVAPNKPSEEEYQRRLRESYTGGFVKEPEKGLWENIVYLDFRALYPSIIITHNVSPDTLNLEGC
KNYDIAPQVGHKFCKDIPGFIPSLLGHLLEERQKIKTKMKETQDPIEKILLDYRQKAIKLLANSFYGYYGYAKARWYCKECAESVT
AWGRKYIELVWKELEEKFGFKVLYIDTDGLYATIPGGESEEIKKKALEFVKYINSKLPGLLELEYEGFYKRGFFVTKKRYAVIDEE
GKVITRGLEIVRRDWSEIAKETQARVLETILKHGDVEEAVRIVKEVIQKLANYEIPPEKLAIYEQITRPLHEYKAIGPHVAVAKKL
AAKGVKIKPGMVIGYIVLRGDGPISNRAILAEEYDPKKHKYDAEYYIENQVLPAVLRILEGFGYRKEDLRYQKTRQVGLTSWLNIK
KSSGazFGSLEHHHHHH

*StuI*

MGSVSAVEQVFLECERARADGDLIQRVSASDKEYHFQNWVQARIEACRLSYDDPGRNTYPDFRLIHHPEGYEVKGLEFPGREADYD
SNSQVPTGNHGGREVFYVFGRYPKAERGVDEYPVVDLVVCHGSFLNADSEYVHKNKSFRGFGSYGDILVRDRKMYVVPTPFALASG
TAGLATLIVPTEFEPQSDTLVQVGELDRTEVDEVIVSYEFNLQTNEMVTHKAPNLNAGKVHSFRAYRSRGAGDSKPVSLAGGRLSG
azFGSLEHHHHHH

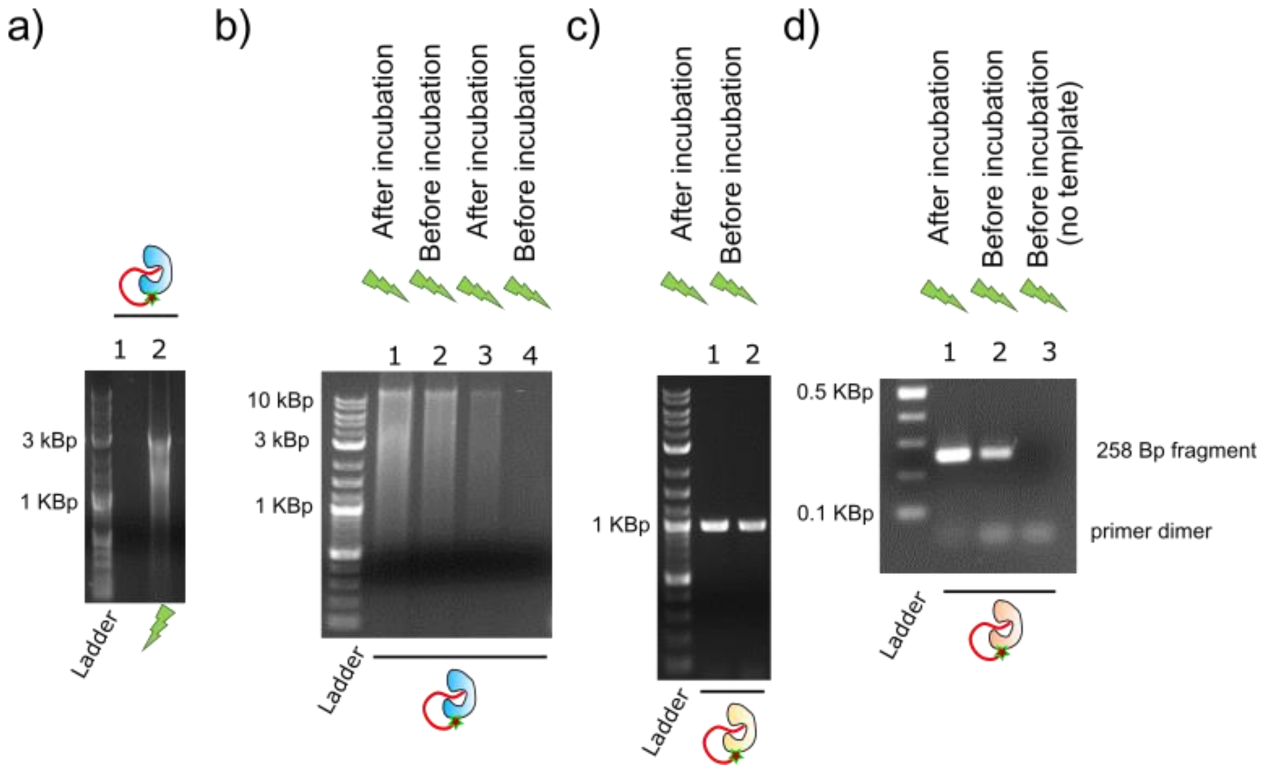


**Figure S1.- Tight-blockage of DNA polymerases and further failure-by-design assays**. **a)** Tight blockage of the activity of Phi pol-PC_oligo. No amplification product was observed when 120 nM of Phi29 pol-PC_oligo was used (lane 1). Activity was recovered after a 10 s light pulse with 315 nm UV (lane 2). **b), c),** and **d)** Independent failure-by-design experiments were performed to corroborate the results shown in Figure 5. **b)** Whole genome amplification by Phi pol-PC_oligo. The hexamers concentration in reactions in lane 1 and 2 was 6.25 µM, and 3.12 µM in lane 3 and 4. **c)** Light-start PCR amplification of Bir A gene with Pfu pol-PC_oligo. 10 nM of enzyme was used and 1 µl of the diluted *E. coli* chromosomic DNA sample. **d)** Ligth-Start with Taq pol-PC_oligo. The same conditions as in Figure 5c were used in this case.

**Figure S2 Fidelity of Taq pol–PC_oligo and Pfu pol-PC_oligo enzymes.** In order to rule out a significant effect on the fidelity of the amplification reaction, we sequenced PCR products amplified by the oligo-modified enzymes. The Bir A gene from *E. coli* (GenBank: M15820.1) was PCR amplified, gel-purified and sent to sequencing (Eurofins Genomics, Germany). The gene was amplified using unmodified Taq pol, unmodified Pfu pol, Tap pol-PC_oligo and Pfu pol-PC_oligo, and the sequences obtained by the unmodified and oligo-modified versions compared (a pulse of 120 s 365 nm UV light was used for the activated enzymes). The first 50 and last 100-200 nucleotides of the sequencing reaction were omitted due to limitations of the sequencing reaction. In none of both cases, differences between the sequences retrieved by the unmodified enzymes and the oligo-modified ones were detected (see sequence alignment below). Furthermore, the sequences show 100% identity with the Bir A deposited sequence (GenBank: M15820.1). Altogether, the results are consistent with a conserved fidelity of the light-activated reactions.

**Pfu pol *vs* Pfu pol-PC_oligo**

```
Alignment of Sequence_1:  [SeqBirA_Pfu.txt.xdna] with  Sequence_2:
[SeqBirA_Pfu_light.txt.xdna]

Similarity : 849/849 (100.00 %)

Seq_1  1     AAGCCCCCTGTTTGTCTATTCCGCGTGAAATGCCAAATATTTCTTTATCACCAATGATAA  60
             ||||||||||||||||||||||||||||||||||||||||||||||||||||||||||||
Seq_2  1     AAGCCCCCTGTTTGTCTATTCCGCGTGAAATGCCAAATATTTCTTTATCACCAATGATAA  60


Seq_1  61    GTTTCACTGGGCGATTAATAAAATTATCCAGCTTTTCCCAGCGCGACAGATAAGGTGCCA  120
             ||||||||||||||||||||||||||||||||||||||||||||||||||||||||||||
Seq_2  61    GTTTCACTGGGCGATTAATAAAATTATCCAGCTTTTCCCAGCGCGACAGATAAGGTGCCA  120


Seq_1  121   ATCCTTCTTGTTCGAAGAGTTCCAACGCAGCACGTAATTCACGTATTAGCATGGCCGCCA  180
             ||||||||||||||||||||||||||||||||||||||||||||||||||||||||||||
Seq_2  121   ATCCTTCTTGTTCGAAGAGTTCCAACGCAGCACGTAATTCACGTATTAGCATGGCCGCCA  180


Seq_1  181   ACGTATTACGATCGAGATTGATCCCCGCTTCCTGCAGCGTGATCCACCCCTGATTAACGA  240
             ||||||||||||||||||||||||||||||||||||||||||||||||||||||||||||
Seq_2  181   ACGTATTACGATCGAGATTGATCCCCGCTTCCTGCAGCGTGATCCACCCCTGATTAACGA  240


Seq_1  241   CACTCTCTTCAACACGGCGCATTGCCATGTTGATCCCGGCTCCAATGACTATTTGCGCCG  300
             ||||||||||||||||||||||||||||||||||||||||||||||||||||||||||||
Seq_2  241   CACTCTCTTCAACACGGCGCATTGCCATGTTGATCCCGGCTCCAATGACTATTTGCGCCG  300


Seq_1  301   CATCGCCAGTTTTGCCAGTCAGCTCCACCAGAATGCCTGCCAGCTTGCGATCCTGCAGAT  360
             ||||||||||||||||||||||||||||||||||||||||||||||||||||||||||||
Seq_2  301   CATCGCCAGTTTTGCCAGTCAGCTCCACCAGAATGCCTGCCAGCTTGCGATCCTGCAGAT  360


Seq_1  361   AGAGGTCATTAGGCCATTTAACACGAACTTTATCTGCACCCAGCTTGCGTAATACTTCCG  420
             ||||||||||||||||||||||||||||||||||||||||||||||||||||||||||||
Seq_2  361   AGAGGTCATTAGGCCATTTAACACGAACTTTATCTGCACCCAGCTTGCGTAATACTTCCG  420


Seq_1  421   CCATCACGATACCGATAACCAGACTTAAACCAATCGCCGCCGCCGGGCCTTGTTCCAGAC  480
```

```
               ||||||||||||||||||||||||||||||||||||||||||||||||||||||||||||
Seq_2  421     CCATCACGATACCGATAACCAGACTTAAACCAATCGCCGCCGCCGGGCCTTGTTCCAGAC  480


Seq_1  481     GCCAGAACATCGACAAATATAAGTTTGCGCCAAAAGGCGAAAACCATTTCCGACCCCGGC  540
               ||||||||||||||||||||||||||||||||||||||||||||||||||||||||||||
Seq_2  481     GCCAGAACATCGACAAATATAAGTTTGCGCCAAAAGGCGAAAACCATTTCCGACCCCGGC  540


Seq_1  541     GACCACGGCCAGCCTGCTGGTATTCTGCAATGCAAGCATCGCCCGATTTAAGCTCTCCGA  600
               ||||||||||||||||||||||||||||||||||||||||||||||||||||||||||||
Seq_2  541     GACCACGGCCAGCCTGCTGGTATTCTGCAATGCAAGCATCGCCCGATTTAAGCTCTCCGA  600


Seq_1  601     TACGATCAAGAAGGTACTGATTCGTGGAGTCAATCACTGGCAGCACGGCTACACTACCGC  660
               ||||||||||||||||||||||||||||||||||||||||||||||||||||||||||||
Seq_2  601     TACGATCAAGAAGGTACTGATTCGTGGAGTCAATCACTGGCAGCACGGCTACACTACCGC  660


Seq_1  661     CATCCAGCTGACCCAATATCTGTTTAGCATTAAGTAACTGGATAGGCTCAGGCAGGCTGT  720
               ||||||||||||||||||||||||||||||||||||||||||||||||||||||||||||
Seq_2  661     CATCCAGCTGACCCAATATCTGTTTAGCATTAAGTAACTGGATAGGCTCAGGCAGGCTGT  720


Seq_1  721     ATCCTTTACCCGGAACGGTAAAGACATCAACGCCCCAGTCACGCAGTGTCTGAATGTGTT  780
               ||||||||||||||||||||||||||||||||||||||||||||||||||||||||||||
Seq_2  721     ATCCTTTACCCGGAACGGTAAAGACATCAACGCCCCAGTCACGCAGTGTCTGAATGTGTT  780


Seq_1  781     TATTAATAGCCGCCCGGCTCATTCCCAGCGTTTCACCCAACTGCTCGCCAGAGTGAAATT  840
               ||||||||||||||||||||||||||||||||||||||||||||||||||||||||||||
Seq_2  781     TATTAATAGCCGCCCGGCTCATTCCCAGCGTTTCACCCAACTGCTCGCCAGAGTGAAATT  840


Seq_1  841     CACCGTTCG  849
               |||||||||
Seq_2  841     CACCGTTCG  849
```

**Taq pol vs Taq pol-PC_oligo**

Alignment of Sequence_1:  [SeqBirA_Taq.txt.xdna] with  Sequence_2:
[SeqBirA_Taq_light.txt.xdna]

Similarity : 750/750 (100.00 %)

```
Seq_1  1       AAGCCCCCTGTTTGTCTATTCCGCGTGAAATGCCAAATATTTCTTTATCACCAATGATAA  60
               ||||||||||||||||||||||||||||||||||||||||||||||||||||||||||||
Seq_2  1       AAGCCCCCTGTTTGTCTATTCCGCGTGAAATGCCAAATATTTCTTTATCACCAATGATAA  60


Seq_1  61      GTTTCACTGGGCGATTAATAAAATTATCCAGCTTTTCCCAGCGCGACAGATAAGGTGCCA  120
               ||||||||||||||||||||||||||||||||||||||||||||||||||||||||||||
Seq_2  61      GTTTCACTGGGCGATTAATAAAATTATCCAGCTTTTCCCAGCGCGACAGATAAGGTGCCA  120


Seq_1  121     ATCCTTCTTGTTCGAAGAGTTCCAACGCAGCACGTAATTCACGTATTAGCATGGCCGCCA  180
               ||||||||||||||||||||||||||||||||||||||||||||||||||||||||||||
Seq_2  121     ATCCTTCTTGTTCGAAGAGTTCCAACGCAGCACGTAATTCACGTATTAGCATGGCCGCCA  180


Seq_1  181     ACGTATTACGATCGAGATTGATCCCCGCTTCCTGCAGCGTGATCCACCCCTGATTAACGA  240
               ||||||||||||||||||||||||||||||||||||||||||||||||||||||||||||
```

```
Seq_2  181    ACGTATTACGATCGAGATTGATCCCCGCTTCCTGCAGCGTGATCCACCCCTGATTAACGA  240

Seq_1  241    CACTCTCTTCAACACGGCGCATTGCCATGTTGATCCCGGCTCCAATGACTATTTGCGCCG  300
              ||||||||||||||||||||||||||||||||||||||||||||||||||||||||||||
Seq_2  241    CACTCTCTTCAACACGGCGCATTGCCATGTTGATCCCGGCTCCAATGACTATTTGCGCCG  300

Seq_1  301    CATCGCCAGTTTTGCCAGTCAGCTCCACCAGAATGCCTGCCAGCTTGCGATCCTGCAGAT  360
              ||||||||||||||||||||||||||||||||||||||||||||||||||||||||||||
Seq_2  301    CATCGCCAGTTTTGCCAGTCAGCTCCACCAGAATGCCTGCCAGCTTGCGATCCTGCAGAT  360

Seq_1  361    AGAGGTCATTAGGCCATTTAACACGAACTTTATCTGCACCCAGCTTGCGTAATACTTCCG  420
              ||||||||||||||||||||||||||||||||||||||||||||||||||||||||||||
Seq_2  361    AGAGGTCATTAGGCCATTTAACACGAACTTTATCTGCACCCAGCTTGCGTAATACTTCCG  420

Seq_1  421    CCATCACGATACCGATAACCAGACTTAAACCAATCGCCGCCGCCGGGCCTTGTTCCAGAC  480
              ||||||||||||||||||||||||||||||||||||||||||||||||||||||||||||
Seq_2  421    CCATCACGATACCGATAACCAGACTTAAACCAATCGCCGCCGCCGGGCCTTGTTCCAGAC  480

Seq_1  481    GCCAGAACATCGACAAATATAAGTTTGCGCCAAAAGGCGAAAACCATTTCCGACCCCGGC  540
              ||||||||||||||||||||||||||||||||||||||||||||||||||||||||||||
Seq_2  481    GCCAGAACATCGACAAATATAAGTTTGCGCCAAAAGGCGAAAACCATTTCCGACCCCGGC  540

Seq_1  541    GACCACGGCCAGCCTGCTGGTATTCTGCAATGCAAGCATCGCCCGATTTAAGCTCTCCGA  600
              ||||||||||||||||||||||||||||||||||||||||||||||||||||||||||||
Seq_2  541    GACCACGGCCAGCCTGCTGGTATTCTGCAATGCAAGCATCGCCCGATTTAAGCTCTCCGA  600

Seq_1  601    TACGATCAAGAAGGTACTGATTCGTGGAGTCAATCACTGGCAGCACGGCTACACTACCGC  660
              ||||||||||||||||||||||||||||||||||||||||||||||||||||||||||||
Seq_2  601    TACGATCAAGAAGGTACTGATTCGTGGAGTCAATCACTGGCAGCACGGCTACACTACCGC  660

Seq_1  661    CATCCAGCTGACCCAATATCTGTTTAGCATTAAGTAACTGGATAGGCTCAGGCAGGCTGT  720
              ||||||||||||||||||||||||||||||||||||||||||||||||||||||||||||
Seq_2  661    CATCCAGCTGACCCAATATCTGTTTAGCATTAAGTAACTGGATAGGCTCAGGCAGGCTGT  720

Seq_1  721    ATCCTTTACCCGGAACGGTAAAGACATCAA  750
              ||||||||||||||||||||||||||||||
Seq_2  721    ATCCTTTACCCGGAACGGTAAAGACATCAA  750
```

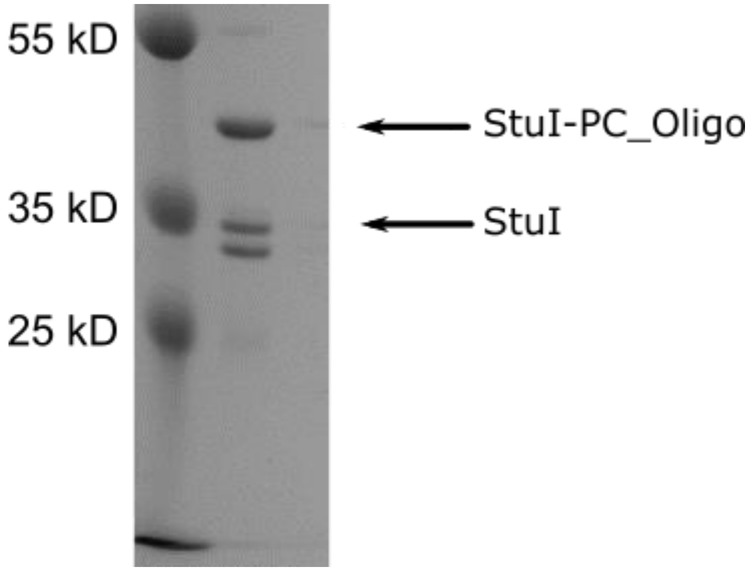


**Figure S3. Sample purity of the StuI-PC_Oligo sample.** SDS-PAGE gel of StuI-PC_oligo sample. Unmodified StuI enzyme was always co-eluted with the StuI-PC_Oligo species. Furthermore, a contaminant of lower molecular weight than StuI co-eluted as well with the enzyme during the previous purification steps. We assigned this contamination to a partially degraded form of the enzyme, as it always co-eluted with the protein after different chromatographic steps (including Nickel affinity purification, cationic exchange, anionic exchange and hydrophobic interaction chromatography). We interpreted the consistent co-elution of the different species to the formation of oligomers, which is consistent with the oligomeric nature of type II restriction enzymes.
